# Characterization and repurposing of the endogenous Type I-F CRISPR-Cas system of *Zymomonas mobilis* for genome engineering

**DOI:** 10.1101/576355

**Authors:** Yanli Zheng, Jiamei Han, Wenyao Liang, Runxia Li, Xiaoyun Hu, Baiyang Wang, Wei Shen, Xiangdong Ma, Lixin Ma, Li Yi, Shihui Yang, Wenfang Peng

## Abstract

Establishment of production platform organisms through prokaryotic engineering represents an efficient means to generate alternatives for yielding renewable biochemicals and biofuels from sustainable resources. *Zymomonas mobilis*, a natural facultative anaerobic ethanologen, possesses many attractive physiological attributes, making it an important industrial microorganism. To facilitate the broad applications of this strain for biorefinery, an efficient genome engineering toolkit for *Z. mobilis* was established in this study by repurposing the endogenous Type I-F CRISPR-Cas system upon its functional characterization, and further updated. This toolkit includes a series of genome engineering plasmids, each carrying an artificial self-targeting CRISPR and a donor DNA for the recovery of recombinants. Using the updated toolkit, genome engineering purposes were achieved with efficiencies of up to 100%, including knockout of *cas3* gene, replacement of *cas3* with the mCherry-encoding *rfp* gene, nucleotide substitutions in *cas3*, and deletion of two large genomic fragments up to 10 kb. This study established thus far the most efficient, straightforward and convenient genome engineering toolkit for *Z. mobilis*, and laid a foundation for further native CRISPRi studies in *Z.* mobilis, which extended the application scope of CRISPR-based technologies, and could also be applied to other industrial microorganisms with unexploited endogenous CRISPR-Cas systems.

## INTRODUCTION

Nearly 50% of bacteria and 90% of archaea equip themselves with adaptive immune systems to defend against invading genetic elements such as phages and plasmids (1). The immunity is CRISPR RNA (crRNA)-based and Cas-driven, functioning in three distinct molecular steps. The Cas proteins encoded by *cas* genes help the integration of short DNA stretches (spacers) into the CRISPR array in a polarized manner (spacer adaptation), process the CRISPR transcript into mature crRNAs (crRNA biogenesis), and execute the crRNA-guided target DNA/RNA destruction (target interference) (2,3). During the interference, targeting a crRNA-matching sequence (protospacer) in an attacking DNA rather than the corresponding spacer stored in the genomic CRISPR bank is dependent on a conserved short sequence immediately flanking the protospacer defined as PAM (protospacer adjacent motif) (4). Additionally, the interference also relies on a seed sequence which is generally the first 8-12 nucleotides following the PAM (5,6).

Various CRISPR-Cas systems have been grouped into six main types (Type I to VI) and basically employ two classes of effector complexes to achieve interference. Class 1 systems involving Type I, III and IV encode multi-subunit effector complexes; while Class 2 systems involving Type II, V and VI have single-Cas ones (7,8). Given the simplicity, the DNA-targeting effectors from Class 2, *i.e.* Type II CRISPR-Cas9 and Type V CRISPR-Cpf1, were developed as genome engineering tools right after their discovery and characterization and have now been widely applied in both eukaryotes and prokaryotes (9-14). In particular, they have enabled rapid, easy yet accurate construction of strains for some production platform microorganisms, *e.g. Saccharomyces cerevisiae* and *Escherichia coli* (15-18), contributing significantly to the bio-based economy. For the bio-based economy, other alternative production microorganisms are also essentially required to generate sustainable alternatives for yielding green chemicals and biofuels from environmentally friendly resources, and become increasingly important. Unfortunately, for the most majority of prokaryotes, the Class 2 CRISPR-based machineries are generally hardly transferable, possibly due to their large size and severe toxicity to host cells. This has largely limited their application in many alternative production hosts, especially some industrially important non-model species including *Zymomonas mobilis*.

*Z. mobilis* is a natural facultative anaerobic ethanologen for the production of cellulosic biofuels and possesses many attractive physiological attributes. For instance, *Z. mobilis* is generally regarded as safe (GRAS), and is capable of tolerating a high ethanol concentration up to 16% (*v/v*) and a broad pH range (3.5-7.5) for ethanol production. Uniquely, *Z. mobilis* takes the Entner-Doudoroff (ED) pathway to anaerobically ferment glucose for ethanol generation which was, interestingly, found in strict aerobic microorganisms (19). Glycolysis via the ED pathway in *Z. mobilis* exhibits several advantages over that through the classical Embden-Meyerhof-Parnas (EMP) pathway in other model microbes such as *S. cerevisiae* and *E. coli* (20-25). Firstly, during the fermentation no advanced aeration control is required, which thus reduces the production cost. Secondly, the fermentation produces relatively 50% less of ATP, giving an improved ethanol yield. Furthermore, as *Z. mobilis* has a high-specific cell surface, it uses glucose with a higher uptake rate than *S. cerevisiae* and *E. coli* do, leading to greater ethanol productivity. These, together with other desirable characteristics, make *Z. mobilis* a robust workhorse candidate for biorefinery practices.

Since the complete genome sequence and functional re-annotation were reported and updated (26-28), in order to fully take the advantages of *Z. mobilis*’ capabilities in biorefinery, a series of genetic tools, including shutter vectors and transformation methods, have been explored and now routinely used (20,29,30). These significant efforts have laid solid foundation for conducting genetic manipulations in *Z. mobilis*. However, these routine genetic manipulation tools are apparently not sufficient for the rapid development of metabolic engineering in the synthetic biology era, which requires high-throughput genome engineering tools, *e.g.* the CRISPR-based toolkits, that have not been completely developed for *Z. mobilis*.

In recent years, several endogenous CRISPR-Cas systems, with the emphasis on Type I, have been harnessed as an alternative strategy to replace the Class 2 CRISPR-based toolkits for prokaryotic engineering in several archaea and bacteria (31-36). Notably, an endogenous Type I-B system was developed as an efficient genome editing toolkit for *Clostridium tyrobutyricum*, an important industrial microorganism for acetone-butanol-ethanol (ABE) production. Using the toolkit, *C. tyrobutyricum* was engineered for n-butanol production and the engineered strains produced butanol with the titer of as historically record high as 26.2 g/L in a batch fermentation (35). However, to the best of our knowledge, this represents thus far the only one native CRISPR-based toolkit for an industrial non-model microorganism. With the growing global demand for alternative sustainable biofuels and biochemicals, more such powerful toolkits for many other industrially important microorganisms are certainly emergently required.

In this study, we characterized the DNA interference capability of the Type I-F CRISPR-Cas system in the type strain *Z. mobilis* ZM4. Upon the characterization, an efficient native Type I-F CRISPR-Cas-based genome editing toolkit was established and with which different genome engineering purposes, including gene deletion/insertion (replacement), point mutation, and large fragment (of up to 10 kb) deletion, were readily achieved. This work provided a versatile and powerful genetic manipulation toolkit for the development and further improvement of *Z. mobilis* as an ideal chassis for biorefinery and synthetic biology studies.

## MATERIALS AND METHODS

### Strains, growth conditions and electroporation transformation of *Z. mobilis*

*Z. mobilis* ZM4 and derivatives constructed in this work are listed in **Table 1**. *Z. mobilis* strains were grown at 30°C in an RMG2 medium (20 g/L glucose, 10 g/L yeast extract, 2 g/L KH_2_PO_4_). If required, spectinomycin was supplemented to a final concentration of 200 µg/mL for *Z. mobilis* and 50 µg/mL for *E. coli*. *Z. mobilis* competent cells were prepared as previously described (30) and transformed with plasmids by electroporation (Bio-Rad Gene Pulser, 0.1-cm gap cuvettes, 1.6 kV, 200 ohms, 25 µF).

**Table 1.** *Zymomonas* strains used in this work

| Strains | Genotype and features | Reference |
| --- | --- | --- |
| ZM4 | *Z. mobilis* subsp. *mobilis* ZM4 (wild-type strain) | (27) |
| $\Delta cas3$ | ZM4 derivative with the entire *cas3* gene deleted | This work |
| $\Delta cas3::rfp$ | ZM4 derivative with the replacement of *cas3* with an *rfp* gene | This work |
| $\Delta Cas$ | ZM4 derivative with the entire *cas* gene cluster deleted | This work |
| $\Delta 10k$ | ZM4 derivative with the deletion of a 10,021-bp genomic fragment | This work |
| $\Delta ZMO0028$ | ZM4 derivative with the entire *ZMO0028* gene deleted | This work |

### Construction of plasmids

Interference plasmids were constructed with two spacers, Spacer7 of CRISPR2 (C2S7) and Spacer4 of CRISPR3 (C3S4). Oligonucleotides were designed to bear the entire sequences of the selected spacers following a 5’-CCC-3’ PAM or the last 3 nucleotides of the repeat (5’-AAA-3’). Each set of oligonucleotides was annealed by heating to 95 °C for 5 minutes and subsequently cooling gradually down to room temperature, giving double-stranded DNA fragments with protruding ends matching DNA ends after double digestion by *XbaI* and *EcoRI*. These DNA fragments were ligated with a linear vector (pEZ15Asp, an *E. coli*-*Z. mobilis* shuttle vector (30)) linearized by *XbaI* and *EcoRI*, yielding the interference plasmids (pInt plasmids) and the corresponding reference plasmids (pRef plasmids) listed in **Table 2**. For the construction of pCas3-Int and pCas3-ref plasmids, a *cas3*-expression cassette was generated by splicing and overlap extension PCR (SOE-PCR) (37) and individually cloned to pInt-C3S4 or pRef-C3S4 plasmid by the T5 exonuclease-dependent DNA assembly (TEDA) method (38).

**Table 2.**
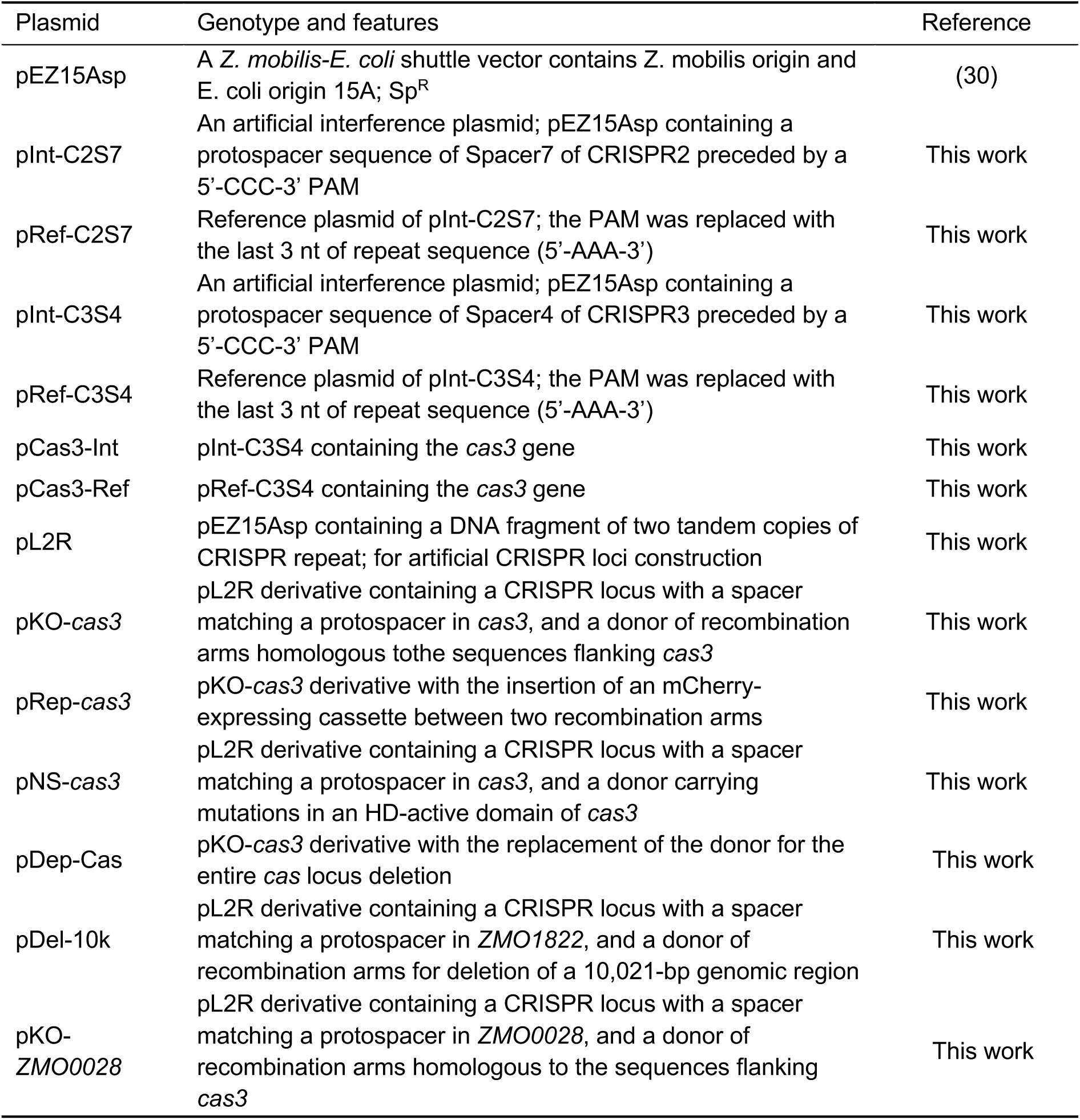
Plasmids used in this work

A number of artificial CRISPR expression plasmids were in play in this study. In order to facilitate the construction, an initial plasmid vector pL2R was constructed based on the pEZ15Asp vector. First, a DNA block composed of the leader sequence of the chromosomal CRISPR2 as a promoter and two CRISPR repeats spaced by two *BsaI* restriction sequences in opposite orientation was synthesized from GenScript (Nanjing, China), and used as a template for PCR amplification with the primer pair of L2R-XbaI-F/L2R-EcoRI-R (**Table 3**). Then, the PCR product was digested with *XbaI* and *EcoRI* and subsequently inserted into the pEZ15Asp vector at the same sites, giving pL2R. Digestion of pL2R with *BsaI* generated a linearized plasmid having protruding repeat sequences of 4 nt at both ends. Double-stranded spacer DNAs were prepared by annealing two spacer oligonucleotides through heating to 95°C for 5 minutes and then cooling gradually down to room temperature. The spacer fragments were designed to carry 4 nt protruding ends complementary to those in the linearized pL2R. Therefore, a number of plasmids each bearing an artificial mini-CRISPR with a self-targeting spacer (pST plasmids, **Table 2**) were yielded by individually ligating the spacer inserts with the linearized pL2R vector. Subsequently, donor DNA fragments each containing a mutant allele of a target gene were generated by SOE-PCR (37) and individually cloned onto their cognate pST plasmids through the TEDA method (38), yielding the all-in-one genomic engineering plasmids listed in **Table 2**. In addition, DNA of the mCherry expression cassette was synthesized from GenScript (Nanjing, China) and used for constructing the donor DNA of the pRep-cas3 plasmid (**Table 2**).

**Table 3.** Transformation efficiencies (TE) and engineering efficiencies (EE) of *Z. mobilis*

| Plasmid | TE (cfu/μg DNA) | | EE [% (engineered/tested)] | |
| --- | --- | --- | --- | --- |
| | ZM4 | ΔZMO0028 | ZM4 | ΔZMO0028 |
| pEZ15Asp | $(3.90 \pm 0.62) \times 10^4$ | $(3.79 \pm 0.23) \times 10^6$ | - | - |
| pKO-ZMO0028 | 297 ± 39 | - | 12.5 (1/8) | - |
| pKO-cas3 | 208 ± 11 | $(2.71 \pm 0.33) \times 10^4$ | 37.5 (3/8) | 100 (8/8) |
| pRep-cas3 | 444 ± 109 | $(1.25 \pm 0.20) \times 10^4$ | 25 (2/8) | 100 (8/8) |
| pNS-cas3 | 41 ± 10 | 93 ± 21 | 7.69 (1/13) | 5.26 (1/19) |
| pDep-Cas | 557 ± 128 | $(1.92 \pm 0.47) \times 10^3$ | 25 (2/8) | 50 (4/8) |
| pDel-10k | 256 ± 44 | $(1.30 \pm 0.25) \times 10^3$ | 12.5 (1/8) | 50 (4/8) |

All oligonucleotides were synthesized from GenScript (Nanjing, China) and listed in **Table 3**. Restriction enzymes and T5 exonuclease were purchased from New England Biolabs (Beijing) Ltd, Beijing, China.

### Construction and screening of mutants

The plasmids used for genome engineering were individually introduced into *Z. mobilis* cells. Electroporated cells were spread on RGM agar plates containing spectinomycin at a final concentration of 200 µg/mL (RGMSp) and incubated at 30°C until colonies were observed. Mutant candidates were checked by colony PCR by using primers listed in **Table 3**. The resulting PCR products were analyzed by agarose gel electrophoresis and DNA sequencing (GenScript Biotech Corp., Nanjing, China).

### Curing of genome engineering plasmids

Cells of genome engineering plasmid transformants were grown up in an RMG2 broth with the supplement of spectinomycin. Then, the cells were spread on an RMG agar plate with spectinomycin to form colonies. A few single colonies were picked and individually suspended in 10 µL of sterilized ddH_2_O, and 1µL of each cell suspension was spotted on an RMG agar plate with or without spectinomycin. Cells from the same suspension grew up on the plate without spectinomycin but not on that with spectinomycin were regarded as those lost the genome engineering plasmid.

### FACS analysis

The protocol used for FACS analysis was modified slightly based on a previous study (39). Briefly, cells were washed with phosphate buffer saline (PBS) twice and then resuspended into PBS to a concentration of 10^7^ cells/mL. Cells were analyzed by flow cytometry using Beckman CytoFLEX FCM (Beckman Coulter, Inc, USA) with the phosphate buffered saline as the sheath fluid. The cells fluorescence of mCherry were excited with the 561 nm and detected with PC5.5 (40,41). Data were processed via FlowJo software (FlowJo, LLC, USA).

## RESULTS

### Characterization of the Type I-F CRISPR-Cas system in *Z. mobilis* ZM4

The complete genome sequence of *Z. mobilis* ZM4 was first published in 2005 (27) with annotation and plasmid sequences updated later (26,28). Based on these data, a CRISPR-Cas system was identified, consisting of four CRISPR arrays (CRISPR1-4) with the same repeat (5’-GTTCACTGCCGCACAGGCAGCTTAGAAA-3’) and a *cas* gene cluster. The CRISPR arrays are located at three different loci in the genome. CRISPR2-4 are on the same strand and CRISPR4 is immediately preceded by CRISPR3, whereas CRISPR1 on the opposite strand. They contain 6, 8, 6 and 1 spacers of 32-34 nt, respectively (**Figure 1A**). Two-thirds of the spacers (14/21) are in length of 32 nt. Interestingly, in the CRISPR-Cas++ database (https://crisprcas.i2bc.paris-saclay.fr), only three of the four CRISPR arrays (*i.e.* CRISPR2-4 here) were collected, missing the CRISPR1. The *cas* genes are organized as an operon of *cas1-cas3-csy1-csy2-csy3-csy4*. The sequence of repeats (**Figure 1B**) and the components and architecture of *cas* genes indicate that this CRISPR-Cas system belongs to Type I-F (42).

**Figure 1.**
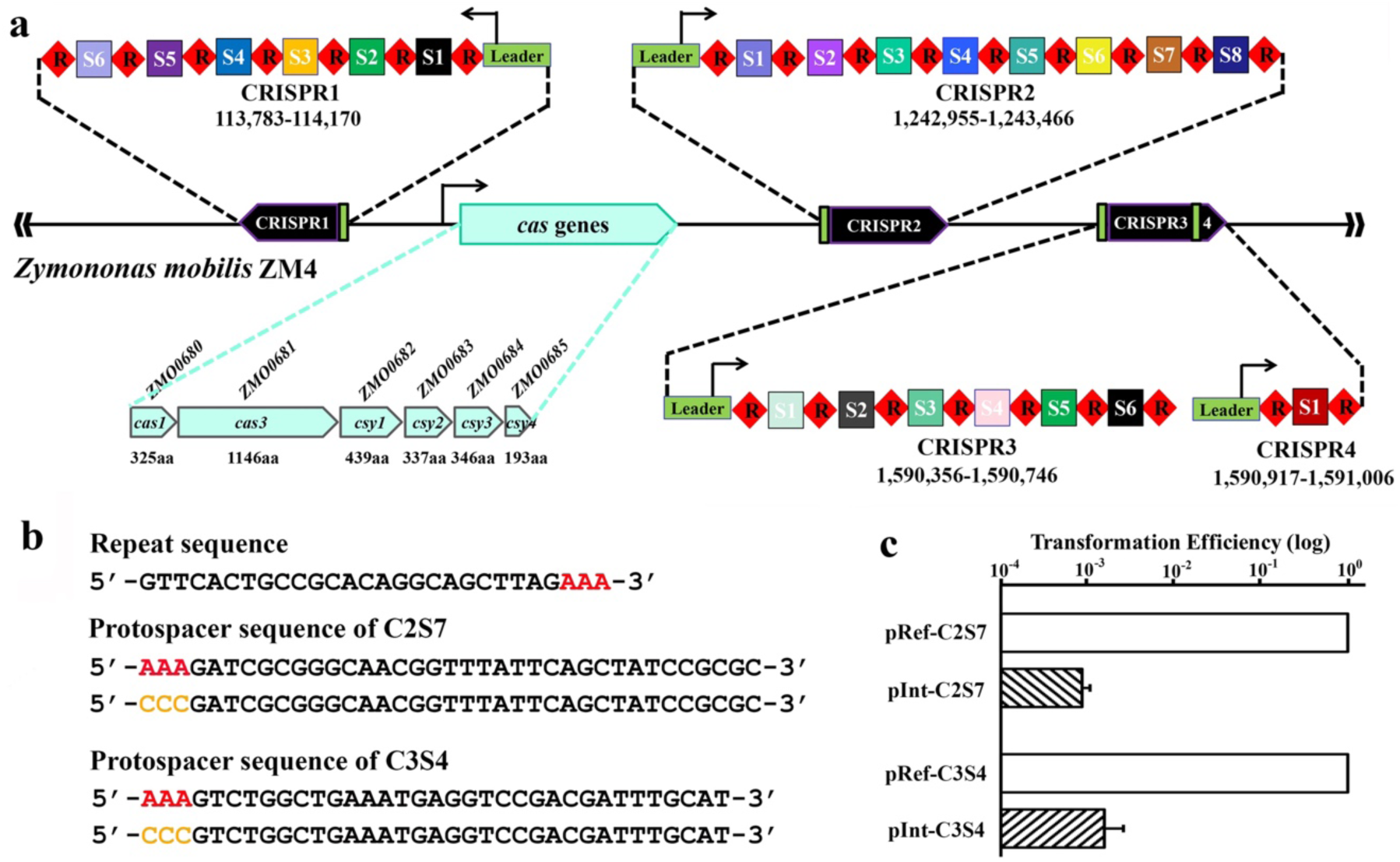
*Z. mobilis* ZM4 CRISPR and *cas* loci and demonstration of the Type I-F DNA interference activity. (**a**) Schematic of CRISPR arrays and a *cas* locus in *Z. mobilis* ZM4. (**b**) Representatives of the repeat sequence, the Spacer7 of CRISPR2 (C2S7) and the Spacer4 of CRISPR3 (C3S4), and the construction of protospacers used in interference plasmids and their corresponding reference plasmids. (**c**) Plasmid interference assay. Two interference plasmids, pInt-C2S7 and pIntC3S4, were used to assess the DNA interference activity of the Type I-F CRISPR-Cas system. Transformation efficiencies of each interference plasmid were expressed as relative values to the efficiencies of their corresponding reference plasmids, the latter of which were set to 1.0.

A prerequisite for exploiting a CRISPR-Cas system for genome engineering is to ensure that the system is functionally active such that self-targeting can be enabled. To test the DNA targeting capability of the Type I-F system of *Z. mobilis* ZM4, we employed a plasmid interference assay since it was shown that a plasmid carrying an artificial protospacer sequence could be efficiently eliminated from the cells expressing the protospacer-matching crRNA (43,44). In addition, for Type I-F systems, a 5’-NCC-3’ sequence was reported to be a functional PAM (45). Hence, we generated protospacer sequences by fusing the 5’-CCC-3’ PAM to the 5’ end of each spacer (**Figure 1B**). Protospacer sequences, of either the 34-bp Spacer7 from CRISPR2 (C2S7) or the 32-bp Spacer4 from CRISPR3 (C3S4), was individually cloned onto the *E. coli*-*Z. mobilis* shuttle vector pEZ15Asp bearing a spectinomycin resistance gene (30), giving the interference plasmid pInt-C2S7 and pInt-C3S4. Meanwhile, in the corresponding reference plasmids (pRef-C2S7 and pRef-C3S4), we replaced the PAM with the last 3 nt of the repeat (5’-AAA-3’), mimicking the genetic structure of CRSIPR arrays in the genome.

We reasoned that the interference plasmids would be recognized and destroyed by the interference complex under the guidance of the corresponding crRNAs produced from the genomic CRISPR loci, leading to no or only few escaped transformants appeared on the agar plates with the supplement of spectinomycin. Instead, the reference plasmids would be considered as a self and thus be ignored by the immune system, allowing many transformants to grow on the same selective plates. As shown in **Figure 1C**, indeed, upon individually introducing the plasmid constructs into *Z. mobilis* cells, hundreds-fold lower transformation rates were obtained with pInt-C2S7 or pInt-C3S4 compared with the reference plasmids, reflecting that the interference plasmids triggered very strong defense response. These results confirmed that the 5’-CCC-3’ motif is a functional PAM and more importantly demonstrated the DNA targeting activity of the Type I-F CRISPR-Cas system, which has the potential to be hijacked for genome engineering in *Z. mobilis*.

### Establishment of the Type I-F CRISPR-Cas-based genome engineering toolkit for *Z. mobilis* ZM4

Given the fact that the endogenous Type I-F CRISPR-Cas system of *Z. mobilis* ZM4 exhibited strong interference activity against the protospacer-bearing plasmids, we were interested in redirecting its DNA cleavage activity to a PAM-flanking sequence on the chromosome for self-targeting and subsequent genome engineering. To this end, self-targeting plasmids (pST plasmids) were constructed to individually carry an artificial CRISPR expression block of leader-repeat-spacer-repeat (**Figure 2A**). In principle, upon introduction of pST plasmid into the *Z. mobilis* ZM4 cells via electroporation, mature crRNAs produced from the artificial CRISPR array would form an effector complex with the Type I-F Cas proteins and reprogram the Type I-F DNA interference activity to make a double-stranded DNA break within the protospacer. Without a DNA repair template, most of the targeted cells may die due to chromosomal DNA cleavage and degradation, whereas some may survive from the targeting as escapers gained uncontrolled mutations that abolish the cleavage (**Figure 2A**). To increase the recovery of desired genotypes in a controllable fashion, a donor DNA consisting of two homology arms for facilitating homologous recombination can be designed to bear expected mutations.

**Figure 2.**
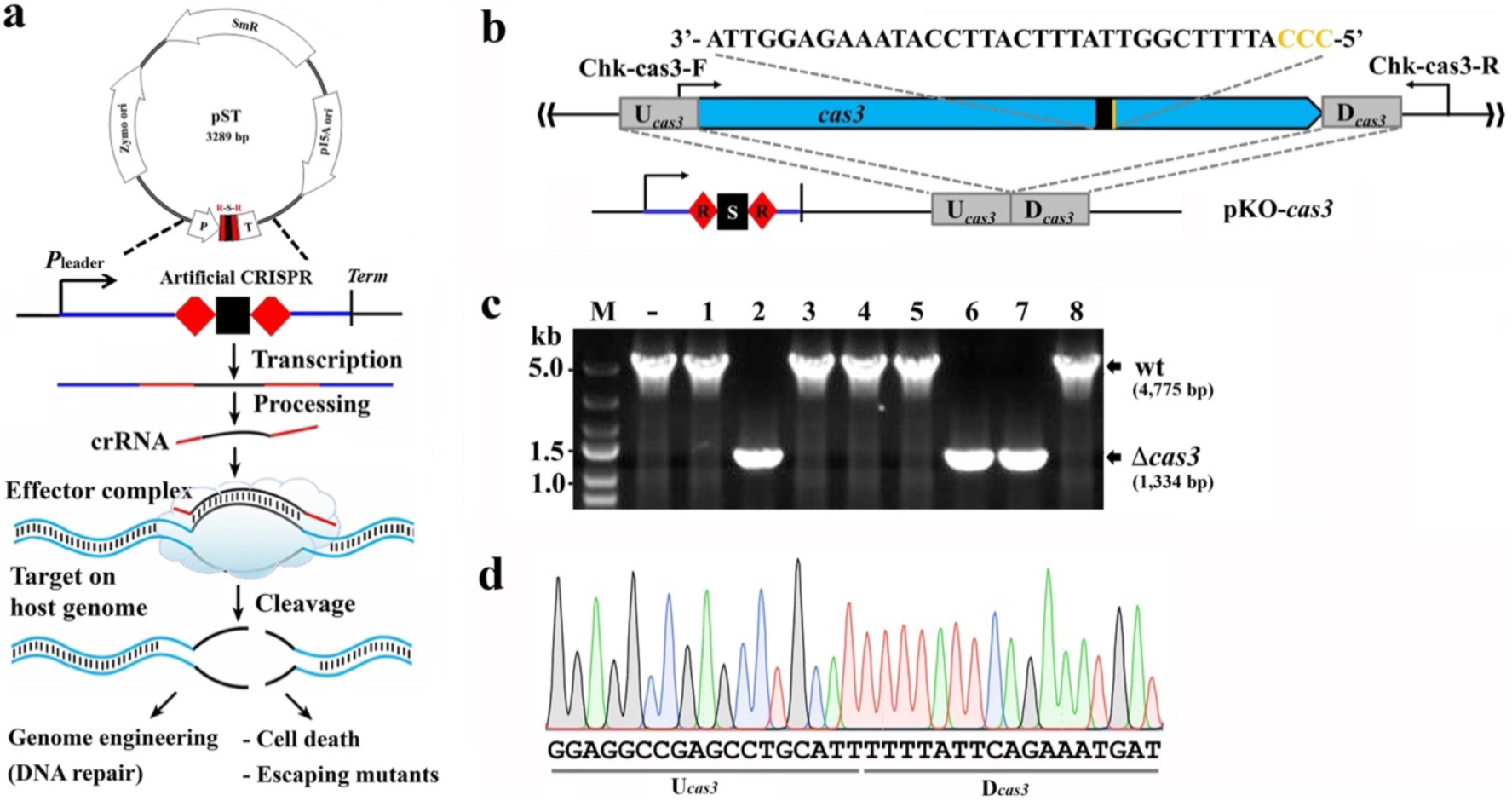
Establishment of the Type I-F CRISPR-Cas-based genome engineering system for *Z. mobilis* ZM4. (**a**) A self-targeting plasmid (pST) contained an artificial CRISPR locus. Self-targeting crRNAs were to be produced from the artificial CRISPR and guided the effector complex to make a double-stranded DNA break in the host genome. The targeted cells either died due to chromosomal DNA degradation, or survived as escaping mutants. (b) Design of the self-targeting CRISPR and the donor DNA in pKO-cas3 for cas3 knockout. The sequence of a 32-nt spacer was shown and the 5’-CCC-3’ PAM was highlighted in orange. (c) Colony PCR screening of cas3 deletion mutant. Primer set of Chk-cas3-F/Chk-cas3-R was used. Predicted sizes of PCR products in wild-type (wt) and the expected δcas3 were indicated with black arrows. -, PCR amplification using genomic DNA of *Z. mobilis* ZM4 as a DNA template. M, DNA size marker. (**d**) Representative chromatographs of Sanger sequencing results of the *cas3* deletion.

The *cas3* (*ZMO0681*) gene was chosen to be engineered for assessing the genome engineering capability of the Type I-F system. The genome engineering plasmid for *cas3* knockout, pKO*-cas3*, was constructed, carrying an artificial *cas3*-targeting CRISPR and the donor DNA. This plasmid was then transformed into *Z. mobilis* ZM4 cells by electroporation. A total of twenty-nine transformants were obtained from three independent transformation events with an average transformation efficiency of 208 ± 11 CFU/µg plasmid DNA (**Table 3**). This transformation rate is hundreds-fold lower than that with the cloning vector pEZ15Asp, indicated that the *cas3*-targeting crNRAs were correctly produced from the plasmid-borne artificial CRISPR locus and had directed efficient self-targeting by the endogenous Type I-F CRISPR-Cas system. Eight of the twenty-nine colonies were analyzed by colony PCR using the primer set of Chk-cas3-F/Chk-cas3-R to select *cas3* knockouts (**Figure 2B**). As shown in **Figure 2C**, a smaller PCR product with a predicted size of 1,334 bp were present in three of the tested strains, *i.e.* Strain 2, 6 and 7, indicating that they were *cas3* deletion mutants. Sequencing results of the PCR products (**Figure 2D**) showed that all these three strains harbored the designed *cas3* deletion while the remaining five strains carried the wild-type *cas3* allele, representing possibly the escaping mutants. Taken together, these results demonstrated that the endogenous Type I-F CRISPR-Cas system could be repurposed for accurate genome engineering with an efficiency of 37.5% (3/8) in *Z. mobilis*.

### Gene replacement by repurposing the endogenous Type I-F system

Metabolic engineering of an alternative production microorganism for generating sustainable alternatives usually requires to introduce heterologous gene or genes of a pathway while eliminate competing endogenous ones. The most convenient and straightforward way to achieve so should be replacing the latter with the former in one step. Gene replacement assay was therefore carried out to examine such capability of the Type I-F system in *Z. mobilis* ZM4. Again, the *cas3* gene was selected as a target to be replaced with the exogenous *rfp* gene encoding the mCherry red fluorescent protein. In this assay, the *cas3*-replacing plasmid, pRep-*cas3*, differed from the pKO-*cas3* by only the donor DNA, where an additional mCherry-expressing block of 1,020 bp was inserted in between of the same two homologous arms as in pKO-*cas3* (**Figure 3A**). Transformation of pRep-*cas3* with *Z. mobilis* ZM4 cells was conducted and transformants were obtained for the following analyses.

**Figure 3.**
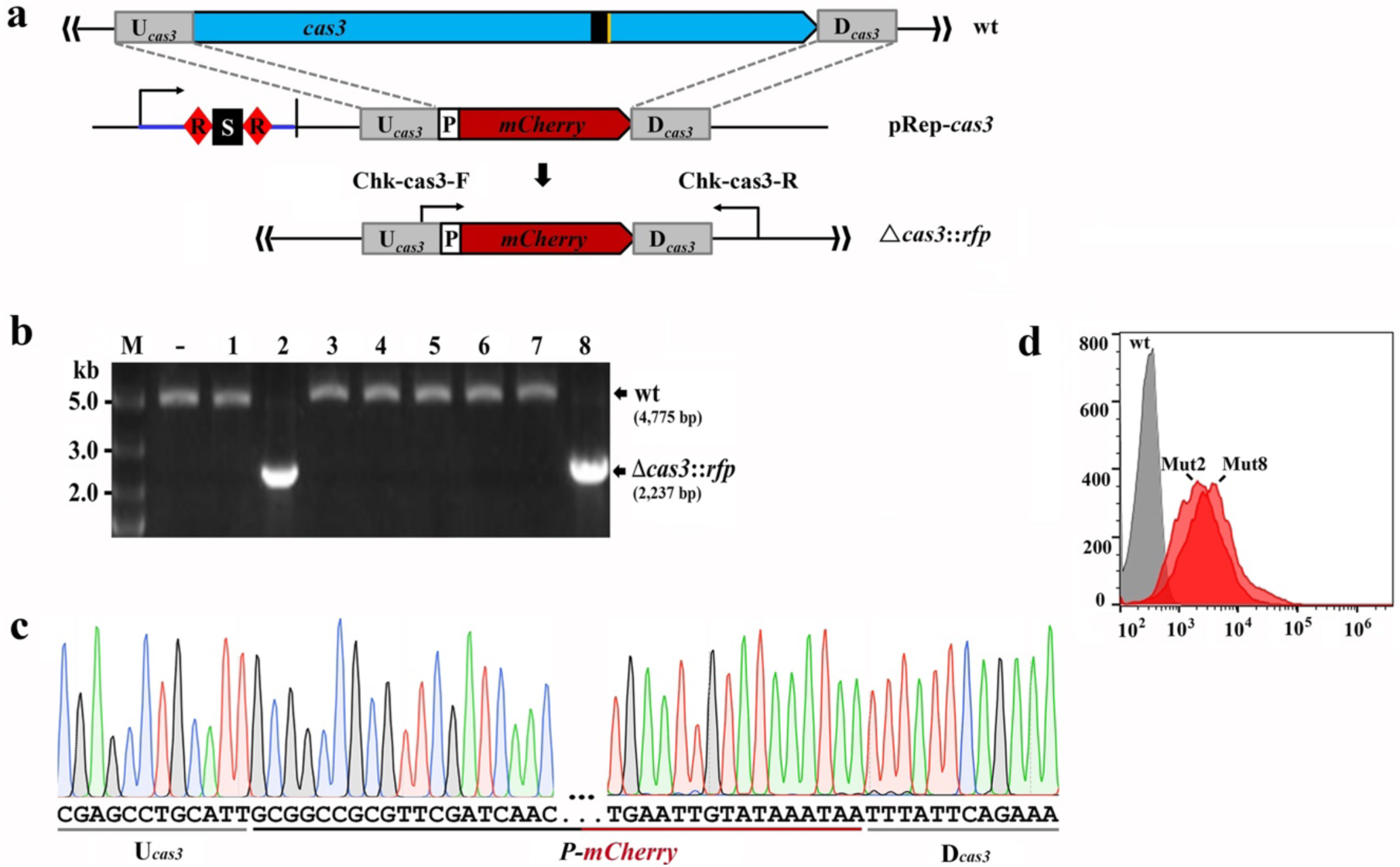
Type I-F CRISPR-mediated replacement of *cas3* with the rfp gene. (**a**) Schematic of *cas3* replacement strategy. pRep-*cas3* carried an additional P-mCherry block in the donor comparing to pKO-*cas3* shown in **Figure 2b**. (**b**) PCR screening of *δcas3∷rfp* recombinants. Predicted sizes of PCR products in wild-type (wt) and the expected *δcas3∷rfp* were indicated with black arrows. -, PCR amplification of genomic DNA of *Z. mobilis* ZM4. M, DNA size marker. (**d**) Signals of mCherry was detected in two Δ*cas3*∷*rfp* mutants (Mut2 and Mut8) whereas not in the wild-type strain (wt) by flow cytometry.

Colony PCR was done using the primer set of Chk-cas3-F/Chk-cas3-R to amplify a DNA fragment encompassing the engineered region from the transformants. A smaller PCR product of the predicted 2,454 bp was yielded in two out of eight tested colonies, representing the Δ*cas3*∷*rfp* mutants (**Figure 3B**), which were further confirmed to be true by DNA sequencing of the PCR products (**Figure 3C**). Moreover, signal of red fluorescent at the same level was detected by flow cytometry in both the edited cells (Mut2 and Mut8), indicating that the heterologously integrated mCherry red fluorescent protein was stably expressed. By contrast, the signal was not detectable in the remaining six strains, which was the same as observed for the wild-type strain (wt) (**Figure 3D**). These results suggested that the Type I-F CRISPR-based toolkit is very useful for precisely integrating heterologous gene or pathway into the host genome for stable production of certain sustainable alternative in *Z. mobilis*.

### *In situ* nucleotide substitutions by repurposing the endogenous Type I-F system

Site-directed mutagenesis is a useful method to study gene or protein functions. Canonically, genes of interest were constructed to contain point mutations *in vitro* and expressed in cells via plasmid (46). This strategy usually changed the copy number and hence the expression level of the target gene. In addition, another approach is to replace the single-nucleotide polymorphisms (SNPs) using homologous recombination containing a selective marker (47-49), which is time consuming or adding heterologous antibiotics gene into the chromosome. In many cases, point mutagenesis was required to be completed in an *in situ* manner. Therefore, we attempted to repurpose the Type I-F system to perform *in situ* nucleotide substitution in *Z. mobilis*.

Once again, the *cas3* gene was used as an engineering target. Cas3 contains nuclease and helicase domains responsible for DNA cleavage in Type I systems (50,51). Here we decided to introduce substitution mutations at an HD-active site of the Cas3 present in the Type I-F system of *Z. mobilis*, ZmCas3, to study its DNA cleavage function. A previous study suggested an HD-active site in the Cas3 of the Type I-F system of *Pseudomonas aeruginosa* (PaCas3), including two Histidine residues located at positions 220 and 221 (H220H221) (52). Alignment of the two Cas3 proteins showed that they shared 42.05% amino acid sequence identity, and conserved Histidine residues corresponding to the H220H221 motif of PaCas3 are present at positions 221 and 222 (H221H222) of ZmCas3 (**Figure 4A**). By carefully inspecting the coding sequences in the vicinity of the H221H222 motif, we found a 5-CCC-3’ PAM and therefore its downstream 32 nt sequence was regarded as a protospacer. Meanwhile, a 1,385-bp donor DNA fragment was yielded by SOE-PCR containing amino acid substitutions of H221A and H222A. Eventually, the genome engineering plasmid for nucleotide substitutions of ZmCas3, pNS-*cas3*, was constructed to carry both the above elements (**Figure 4B**). Transformation with pNS-*cas3* yielded a transformation rate of 41 ± 8 CFU/µg plasmid DNA, which was about one-fifth of that with the pKO-*cas3* plasmid (**Table 3**). Then, colony PCR and Sanger sequencing analyses revealed that accurate nucleotide substitutions (H221A + H222A) occurred (**Figure 4C**) in only one of all the obtained thirteen colonies, showing a recombination recovery rate of 7.69%.

**Figure 4.**
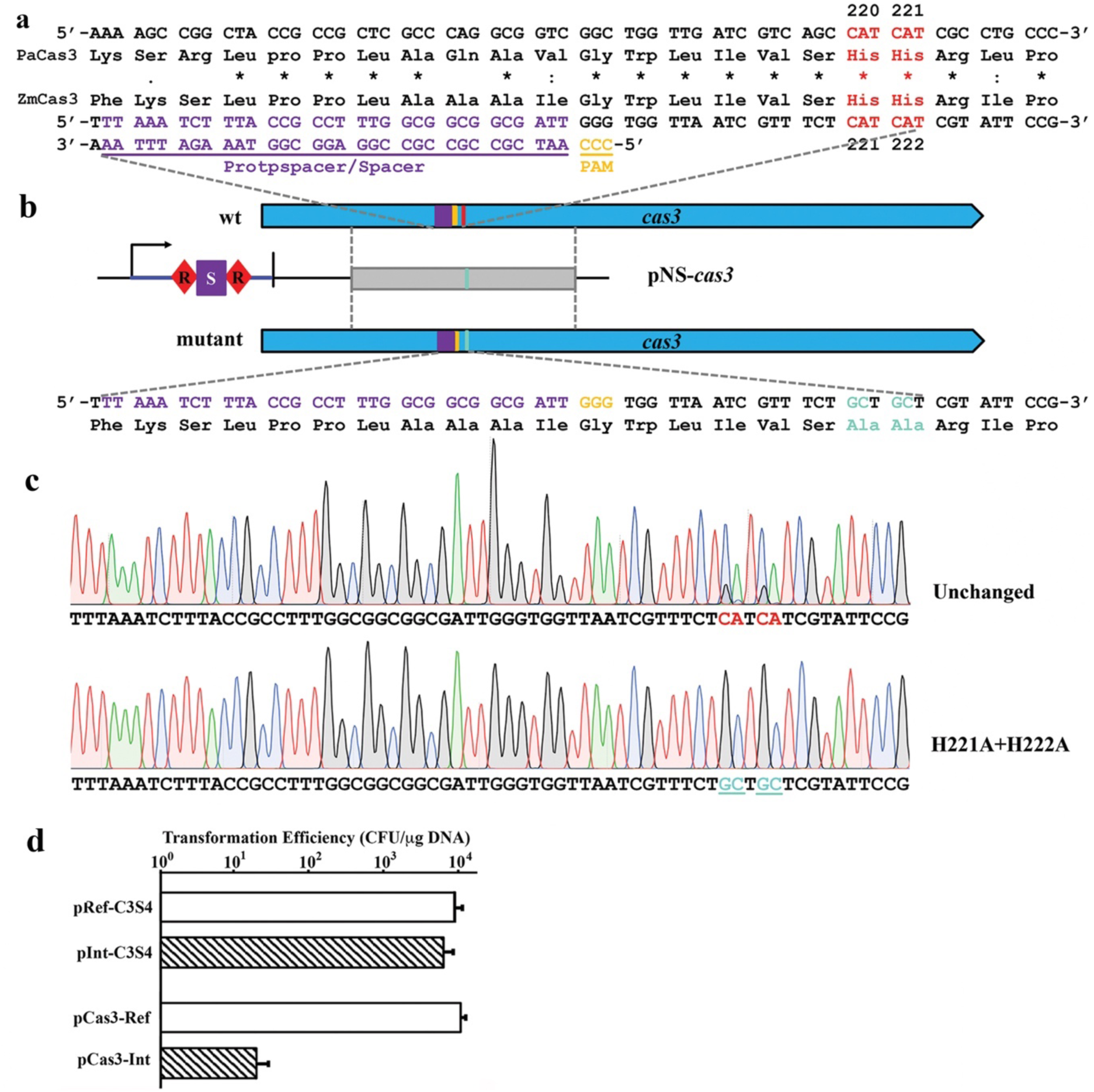
Nucleotide substitutions of an HD-active site of *cas3*. (**a**) Alignment of amino acid sequences containing an HD-active site in the *Z. mobilis* ZM4 Cas3 (*ZMO0681*), ZmCas3, and its *Pseudomonas aeruginosa* homolog, PaCas3. Two conserved amino acids of histidine were selected for constructing alanine substitutions (highlighted in red). (**b**) Schematic showing the mutagenic strategy. The protospacer in *cas3* was indicated in purple while the PAM in orange. Designed mutations in the *cas3* mutant were shown in cyan. (**c**) Representative chromatographs of Sanger sequencing results of the *cas3* mutant. Mutations carried by the mutant were highlighted in cyan, while the corresponding unchanged nucleotides kept by other transformants were shown in red fonts. (**d**) Effect of the nucleotide substitution mutations in *cas3* on plasmid DNA interference was assessed by plasmid interference assay of the *cas3* mutant. pInt-C3S4 and pCas3-Int were used as interference plasmids, while pRef-C3S4 and pCas3-Ref as the corresponding reference plasmids. Three replicates were performed for each.

We found that all the thirteen transformants still contained the pNS-*cas3* plasmid and no any change was found in the artificial CRISPR loci. Moreover, except for the Cas3 mutant, the remaining twelve colonies had no sequence change in the *cas3* alleles. These results suggested that other unwanted mutations in the remaining twelve colonies made the failure of CRISPR immunity. This thus raised a question of whether only the designed substitutions, rather than some other unwanted mutations, had caused the loss of DNA targeting capability of the obtained Cas3 mutant. To address this, the pNS-cas3 plasmid was cured from the Cas3 *mutant* and the cells were used as genetic host for retransformation with interference plasmids.

First, pInt-C3S4 and pRef-C3S4 plasmids were individually retransformed into the Cas3 mutant cells. Tens of thousands transformants appeared with both plasmids, suggestive of no CRISPR immunity in this mutant. Then, two additional plasmids were constructed by individually cloning a *cas3*-expressing cassette on pInt-C3S4 and pRef-C3S4, generating pCas3-Int and pCas3-Ref, respectively, where the expression of Cas3 was under the control of the promoter of the *Z. mobilis adhB* gene encoding the alcohol dehydrogenase II (P_*adhB*_) (53). Expectedly, the Cas3 function would be restored by episomic expression of *cas3* from the plasmids, so that the pCas3-Int plasmid would be targeted again by the Type I-F system. Indeed, no transformant was obtained from the transformation with pCas3-Int in three independent transformations, showing very strong targeting effect on the plasmid-borne artificial protospacer, whereas a very high transformation rate comparable to that with pRef-C3S4 was yielded with pCas3-Ref (**Figure 4D**). These results suggested that the loss of CRISPR immunity in the Cas3 mutant was solely due to the introduced H221A and H222A substitutions, and hence also confirmed the HD-active site for the Cas3 nuclease activity. This turned the Cas3 to be a catalytically deficient Cas3, dCas3, and hence generated a native CRISPR-dCas3 system with great potential for future CRISPRi study.

### Deletion of large genomic fragment by repurposing the endogenous Type I-F system

Minimal microbial genome construction is a straightforward approach for generating a chassis for industrial applications and one means for this purpose is to reduce the genome by deleting the unnecessary genomic fragments (54), including single small-sized genes or large segments. *Z. mobilis* ZM4 has a small genome with the size of only 2.06 Mb (26) that has evolved with a high fraction of essential genes (55), and large fragment deletion could be somehow restricted by the relatively poor performance of the routine genetic manipulation methods for *Z. mobilis* (20). To date, most DNA deletions from the *Z. mobilis* ZM4 genome were small ones, and no large-fragment deletion efforts have been reported. Therefore, having achieved single gene deletion and replacement using the powerful CRISPR-based toolkit established in this study, we were interested in testing the capability of this system in deleting large genomic fragments.

The *cas* gene cluster is an ideal target for this purpose. First, depletion of the Cas proteins had no effect on cell viability. Second, the gene cluster spans a (large) genomic region of 8,446 bp (Figure 5A). Finally, the *cas3* gene is included in the cluster, so that the *cas3*-targeting spacer can be further used here. The genome engineering plasmid for depleting all the Cas proteins (pDep-Cas) in *Z. mobilis* ZM4 contained the same self-targeting CRISPR as that in the pKO-*cas3* plasmid. Transformation with pDep-Cas via electroporation obtained a transformation rate comparable to that obtained in the above experiments. We analyzed the target locus in eight randomly chosen transformants by colony PCR with Chk-Cas-F and Chk-Cas-R primers (**Figure 5A, Supplementary Table 1**). PCR products were produced and subjected to gel electrophoresis, exhibiting bands with two different sizes. The smaller band appeared in two of the colonies, corresponding to the designed *cas* locus deletion mutants of the predicted size of 956 bp (ΔCas) (**Figure 5B**). These PCR products were then sent for DNA sequencing and the results verified that the precise deletion of an 8,446-bp genomic fragment occurred in both colonies (**Figure 5C**). The remaining six colonies should still harbour the *cas* locus as from the genome of which the wild-type band (10,290 bp) was amplified with the same pair of primers (**Figure 5B**).

**Figure 5.**
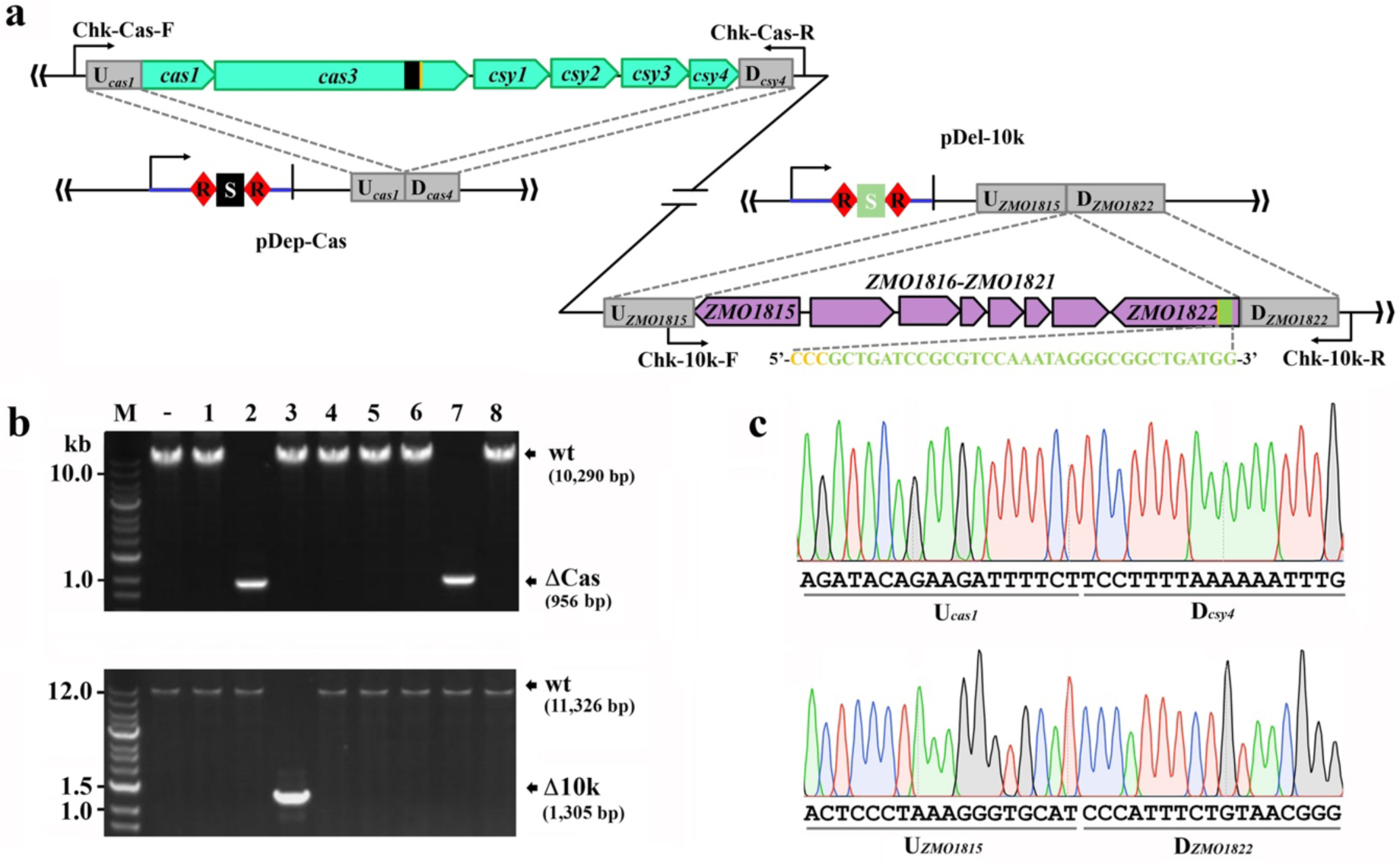
Deletion of large genomic fragments. (**a**) Schematic showing the targeting sites, and the design of self-targeting CRISPRs and the donor DNAs in the corresponding genome engineering plasmids, pDep-Cas and pDel-10k. The CRISPR carried by pDep-Cas is same as that in pKO-*cas3* shown in **Figure 2b**. The sequence of a 32-nt spacer used in pDel-10k was shown and the 5’-CCC-3’ PAM was highlighted in orange. (**b**) PCR screening of mutants with large genomic fragment deletion. Predicted sizes of PCR products in wild-type (wt) and the expected δCas with Chk-Cas-F/Chk-Cas-R and δ10k with Chk-10k-F/Chk-10k-R, respectively, were indicated with black arrows. -, PCR amplification of genomic DNA of *Z. mobilis* ZM4. M, DNA size marker. (**c**) Representative chromatographs of Sanger sequencing results of the entire *cas* gene cluster deletion (upper panel) and the >10 kb deletion (lower panel).

In all the above experiments, the *cas3* gene was taken as or included in a target for engineering. In order to verify that the Type I-F CRISPR-based toolkit can be generally utilized for modification of any gene of interest in *Z. mobilis* ZM4, we attempted to use it to delete another genomic fragment of >10 kb. Based on the genome sequence and functional re-annotation (26), a DNA stretch was considered as a target, containing genes from *ZMO1815* to *ZMO1822* (**Figure 5A**). Among them, the genes of *ZMO1816-1821* was organized in an operon and might play roles in N_2_ fixation as a bypass for the traditional bacterial nitrogen metabolism pathways in *Z. mobilis* isolates (56). The *ZMO1815* and *ZMO1822* genes flanking the operon encode orthologues of TonB-dependent siderophore receptor. Their orthologues in *P. aeruginosa* (PiuA and PirA) were proven to be not essential for cell viability because a *piuA-pirA* double mutant was already obtained (57). In addition, two other genes, *ZMO0188* and *ZMO1631*, in the *Z. mobilis* ZM4 genome also encode orthologues of TonB-dependent siderophore receptor (26) that might play the similar role as *ZMO1815* and *ZMO1822* do. Taken together, *ZMO1816-1821* together with *ZMO1815* and *ZMO1822* are removable and a 10,221-bp genomic region including all of which was eventually designed to be deleted in this work.

A genome engineering plasmid for deleting the fragment of >10k (pDel-10k) was generated by employing the same construction strategy as described in the above experiments (**Figure 5A**). Several transformants were obtained from the transformation with pDel-10k. To uncover the genotype of the transformants, colony CPR was conducted using Chk-10k-F and Chk-10k-R (**Supplementary Table 1**) to amplify a DNA region spanning the engineered locus (**Figure 5A**). PCR products of the predicted size of 1,305 bp were produced in one out of eight analyzed colonies (Δ10k), which were absent from all the rest and the original host (wt) (**Figure 5B**). Finally, sequencing results of the 1305-bp PCR products showed accurate deletion of the 10,021-bp genomic segment as designed (**Figure 5C**).

### Efficiency improvement of the endogenous Type I-F CRISPR-Cas-based genomic engineering toolkit for *Z. mobilis*

Although we have succeeded in various genome engineering purposes with the Type I-F CRISPR-Cas system in *Z. mobilis*, we also noticed that the engineering efficiencies was much lower (6.79-37.5%) than that obtained by repurposing other native Type I systems (31,32,35). This left sufficient room for its further improvement and we therefore sought to update this toolkit to be a more efficient one.

Previous studies showed that bacteria and archaea could escape self-targeting of CRISPR-Cas by generating low-frequency mutations (34,58,59). Consistently, in the plasmid interference assay, when challenged with either the interference plasmid (pInt-C2S4 or pInt-C3S7), always no or only few transformants could be recovered (**Figure 1C**). This indicated that the rate of escaping mutations was generally very low and normally at a fixed level. Inconceivably, such low-frequency mutations even contributed to the obtaining of at least 62.5% of the transformants in this work. This hinted at a low recovery rate of mutants with desired genotype from DNA repair via recombination, and reminded us to pay attention to the plasmid transformation efficiency, because in all the experiments the donor DNAs for facilitating homologous recombination were delivered into the host cells as inserts of the corresponding genome engineering plasmids. The amount of the occurring donor DNAs in a target cell was the principal factor affecting the recombination rate and hence the genome engineering efficiency. Therein, we speculated that the generally low engineering efficiencies yielded in this work were possibly caused by the generally low plasmid transformation efficiency (around 10^4^ CFU/µg DNA) due to the presence of DNA restriction-modification (R-M) systems in *Z. mobilis* (60).

To address the speculation, we decided to deplete the Type IV R-M element ZMO0028 and reassess genome engineering under a background of Δ*ZMO0028*, because inactivation of the ZMO0028 led to an improvement of plasmid transformation efficiency by up to 60-fold (reaching 10^6^ CFU/µg DNA) (60). Deletion mutants of *ZMO0028* were successfully obtained with an editing efficiency of 12.5% (data not shown) by using the genome engineering plasmid, pKO-*ZMO0028* (**Table 2**). Consistent with the previous study (60), in this study, when transformed with the pEZ15Asp vector, the transformation efficiency of Δ*ZMO0028* was about 90-fold higher than that of the wild-type strain, and the same was also true when transformed with the genome engineering plasmids except for pNS-*cas3* (**Table 3**).

Then, Δ*ZMO0028* was used as a genetic host to individually transformed with the genome editing plasmids. Analyses of colony PCR and Sanger sequencing with the transformants obtained from each assay were done as the same did in the above experiments and the results summarized in **Table 3**. Strikingly, under the Δ*ZMO0028* background, substantially enhanced genome engineering rates were obtained for all the genome engineering manipulations. Noteworthily, engineering efficiencies of the knockout of *cas3* and the replacement of *cas3* with *rfp* reached 100%, while that of both the deletions of large genomic fragment improved to 50%. It was seen that the engineering efficiency of the nucleotide substitutions in Cas3 was of 5.26%, even lower than or at a similar level with that obtained under the wild-type background (7.69%). The reason could be that, the pNS-*cas3*-carried donor DNA included the same protospacer as present in the genome and therefore was also targeted by the CRISPR immunity until the inactivation of Cas3 nuclease activity caused by the H221A and H222A substitutions, which will be further discussed in the **DISCUSSION** session below.

## DISCUSSION

In this study, we have established a native CRISPR-Cas-based genome engineering toolkit for the first time for an important industrial microorganism *Z. mobilis*, which overcomes the key restriction of using the Class 2 CRISPR-based genome editing technologies in *Z. mobilis* and in many other prokaryotes and also extends the application scope of CRISPR-based technologies. By using this toolkit, genome engineering options, including gene knockout, insertion or replacement, large genomic fragment deletion, and specific point mutagenesis, can be readily achieved, representing thus far the most efficient and straightforward genome engineering toolkit for *Z. mobilis*. When compared with the routinely used genetic manipulation methods, this CRISPR-base toolkit is much simpler, more convenient and time-saving. For instance, due to the strong selection pressure conferred by CRISPR targeting, no additional selection marker is required. Basically, mutants with designed mutations can be readily obtained in a single round of transformation, spending about only three days, whereas it normally takes at least half a month to achieve so in the routine genetic manipulations. Certainly, the CRISPR-based toolkit established here will largely facilitate further elucidation of the underlying mechanism of inhibitor-tolerant mutants with single-nucleotide polymorphisms (SNPs) (47,48,61), which cannot be easily studied by classical genetics approaches to correlate the robustness phenotypes with their corresponding genotypes. In addition, this toolkit will expedite the development of *Z. mobilis* as a synthetic chassis for sustainable economic biofuel and biochemical productions.

To date, genome engineering with the already developed native Type I CRISPR-based toolkits can be achieved at generally high efficiencies (31,32,35). However, based on the specific genetic backgrounds of various microorganisms, several attempts were made on the optimization of the native CRISPR-based genome engineering toolkits, aiming at high engineering efficiencies. For example, in the studies of repurposing the native Type I-A of *Sulfolobus susfolobus* (31) and Type I-B of *Haloarcula hispanica* (32) the authors first tried providing donor of free linear DNA but failed in obtaining designed mutants or got few at a very low efficiency. Therefore, they inserted the donor into the genome editing plasmid, which was proven to be a very efficient approach to attain advanced engineering efficiencies. Furthermore, Cheng *et al.* also evaluated the effect of the donor DNA lengths on genome engineering efficiency and eventually decided a donor size of 600-800 bp (32). We adopted the same strategy in this study. In another study where the endogenous Type I-B of *Clostridium tyrobutyricum* was exploited for genome engineering, Zhang and co-authors tested the spacer length of the artificial CRISPR on the efficiencies of plasmid transformation and genome engineering (35).

In this study, we demonstrated that the enhancement of plasmid DNA transformation efficiencies through inactivating the Type IV DNA R-M system in *Z. mobilis* led to boosted genome engineering efficiencies of up to 100%, which are comparable to that with other endogenous Type I CRISPR-based prokaryotic engineering toolkits. R-M systems naturally occur in prokaryotes even more broadly than the CRISPR-Cas systems do, pointing towards a fact that both the adaptive CRISPR-Cas immune systems and the innate R-M systems may coexist in many bacteria and archaea (62). Thus, it may explain the perplexity that the diverse CRISPR-Cas systems broadly occur in prokaryotes in nature whereas the native CRISPR-based technologies are much lesser applied. Possibly, in these bacteria and archaea highly efficient native CRISPR-based genome engineering would be only enabled under at least an R-M^-^ background. In addition, other prokaryotic defence modules that also prevent foreign DNA introduction such as the Toxin-Antitoxin (TA) systems would be taken account as well if occurred. Together, the method developed here may serve as an important reference for the development and update of similar toolkits in prokaryotes that harbour active endogenous CRISPR-Cas systems but with low efficiency in the wild-type genetic background.

The reason for low engineering rates of nucleotide substitutions in *cas3* under both wild-type and Δ*ZMO0028* backgrounds could be the essentiality of the PAM position. CRISPR-mediated point mutations were generally exclusively introduced into the PAM or other essential positions in other studies (*e.g.* the seed sequence) to inactivate the DNA interference activity of the system, such that the edited target, as well as the donor DNA provided along with the genome engineering plasmid, would not be persistently targeted by the CRISPR-Cas system. However, in many cases, it is hard to find a protospacer overlapping with the candidate sites and thus suitable for engineering. For example, we could not find any 5’-NCC-3’ PAM-preceded protospacer to make the designed nucleotide substitution positions sit right in the PAM or the seed sequence. Therefore, we had to choose a protospacer close to the engineering sites instead. In this case, until the mutations were introduced to inactivate the Cas3 nuclease activity, both the protospacer sequences in the genome and the donor DNA on the plasmid were cleaved, thus accounting for the low efficiencies yielded for both strains. Nevertheless, the engineering objective was still obtainable by using this Type I-F CRISPR-Cas-based toolkit established here.

In this work, we took the *cas3* gene as a target to study genome engineering in *Z. mobilis*, and different *cas3* mutant strains were successfully obtained. All Type I systems use a complex of Cas proteins to bind to a target while employ Cas3 nuclease to execute protospacer cleavage (63,64). Depletion of Cas3 or keeping a catalytically inactivity Cas3 (dCas3) in the system would still allow the complex to bind to the target DNA but without cutting it, thus converting the system to be a gene repression platform, as the binding complex would act as a transcriptional barrier through preventing RNA polymerase access or transcription elongation depending on the location of the targeted protospacer. Likewise, upon fused with the bacterial RNAP activator SoxS, the dCas3 can be used to recruit RNAP assembly for transcription activation, which is in parallel with the functions of dCas9/dCpf1 in Class 2 systems (65). Indeed, for example, by taking such a strategy, the Type I-B system of *Haloferax volcanii* was harnessed for efficient repression of transcription upon Cas3 depletion. Interestingly, the expression of a catalytically inactive Cas3 via plasmid gave a further enhanced knockdown effect on an endogenous gene (66). Herein, the endogenous Type I-F CRISPR-Cas system could be also developed as CRISPR interference (CRISPRi) and activation (CRISPRa) toolkits (67-69) or similarly as reported recently as CRISPR-assisted multi-dimensional regulation for fine-tuning gene expression (70) and metabolic engineering for *Z. mobilis* and related microorganisms in the future.

---

**Supplementary Table 1.**
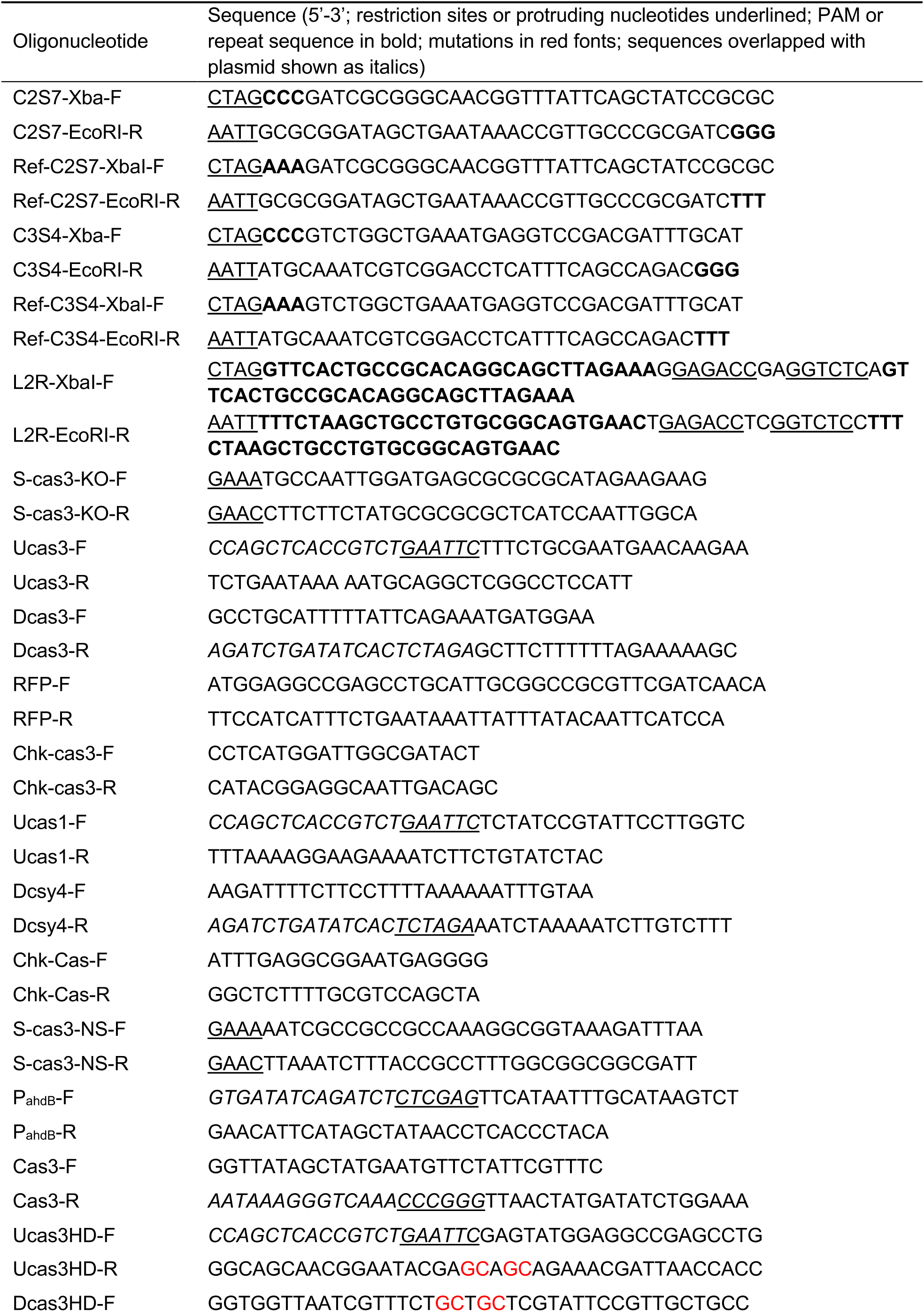
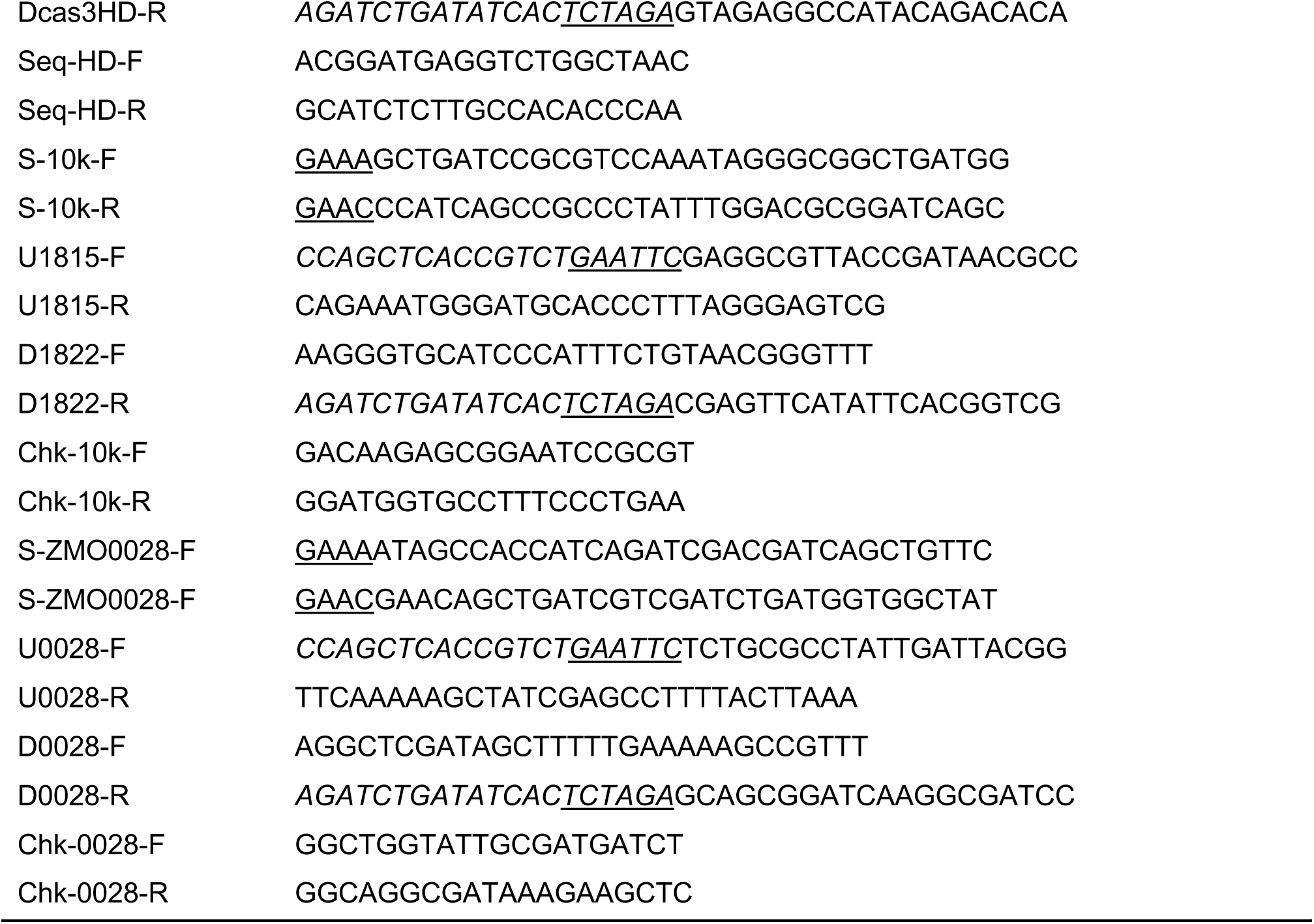
Oligonucleotides used in this work

